# G3BP1 Phosphorylation Regulates Plant Immunity in *Arabidopsis*

**DOI:** 10.1101/2025.05.06.652493

**Authors:** Fatimah Abdulhakim, Aala Abulfaraj, Naganand Rayapuram, Heribert Hirt

**Affiliations:** Darwin21 Desert Research Initiative, Biological and Environmental Science and Engineering Division, 4700 King Abdullah University of Sciences and Technology, 23955-6900 Thuwal, Saudi Arabia; Biological Sciences Department, College of Science & Arts, King Abdulaziz University, Rabigh 21911, Saudi Arabia

## Abstract

Mitogen-activated protein kinase (MAPK) cascades play critical roles in plant immunity by phosphorylating downstream effectors that regulate stress responses. While MAPK-mediated transcriptional regulation has been well examined, the involvement of MAPKs in post-transcriptional and post-translational regulation is still poorly understood. In this study, we identify the RNA binding-protein AtG3BP1 as a phosphorylation target of MPK3, MPK4, and MPK6 and demonstrate that phosphorylation of AtG3BP1 at Ser257 modulates key aspects of *Arabidopsis* immunity. Using phospho-mimic (G3BP1^D^) and phospho-dead (G3BP1^A^) mutants, we investigated the functional consequences of AtG3BP1 phosphorylation. Our data indicate that phosphorylation of AtG3BP1 promotes susceptibility to bacterial infection, suppresses accumulation of reactive oxygen species (ROS), and downregulates salicylic acid (SA) biosynthesis. Furthermore, we demonstrate that AtG3BP1 phosphorylation influences stomatal immunity by maintaining stomatal opening, thereby regulating pre-invasive defense mechanisms. Additionally, we provide evidence that phosphorylation stabilizes AtG3BP1 and prevents its degradation via the proteasome, thus leading to sustained immune signaling. These findings validate AtG3BP1 as a central integrator of MAPK signaling during plant immunity and reveal a new level of post-translational control. This study enhances our understanding of plant defense mechanisms and provides potential targets for engineering disease-resistant crops.

## Introduction

Plants have highly developed immune systems that protect them from pathogens. Plant immune responses are composed of two main pathways: pattern-triggered immunity (PTI) and effector-triggered immunity (ETI) (Bigeard et al., 2015). PTI is activated when pattern recognition receptors (PRRs) detect conserved microbe-associated molecular patterns (MAMPs/PAMPs), such as bacterial flagellin and fungal chitin (Bigeard et al., 2015; Jones & Dangl, 2006). However, successful pathogens secrete effector proteins into plant cells through specialized secretion systems, such as the type III secretion system (T3SS), to suppress PTI and induce effector-triggered susceptibility (ETS) (Macho & Zipfel, 2014; Alhoraibi et al., 2019). In response, plants have evolved intracellular nucleotide-binding leucine-rich repeat (NB-LRR) receptors that recognize pathogen effectors and activate ETI, which often leads to localized cell death to prevent the spread of pathogens (Jones & Dangl, 2006; Dangl & Jones, 2001).

A key component of PTI signaling is the activation of immune mitogen-activated protein kinases (MPKs) MPK3, MPK4, and MPK6. MAPKs are central regulators of plant immune responses (Rodriguez et al., 2010; Meng & Zhang, 2013; Bigeard et al., 2015; Komis et al., 2018) that phosphorylate key downstream proteins to regulate plant defense responses.

To investigate MAPK-mediated phosphorylation dynamics, we conducted an *in vivo* phosphoproteomic study of chromatin-associated proteins from wild-type Col-0 and mapk (*mpk3, mpk4, mpk6*) mutant plants. One-month-old *Arabidopsis* plants were treated for 15 minutes with water or 1 μM flg22, a well-characterized MAMP that activates plant immunity. Among the identified MPK substrates, Ras GTPase-activating protein SH3-domain-binding protein (AtG3BP-1, AT5G48650) was found to be phosphorylated at (Ser/Thr) phospho-(S/T)P sites, which suggests that this protein is a direct substrate of MPK3, MPK4, and MPK6 (Rayapuram et al., 2021). Previous phosphoproteomic studies also identified AtG3BP-1 as a phosphorylated protein, with phosphorylation detected within the DKFGVPAVSLPpSPK sequence, which contains an (S/T)Pmotif, a known MAPK substrate recognition site. Additionally, AtG3BP-1 was categorized as a phosphoprotein with a putative role in RNA metabolism (van Bentem et al., 2006).

AtG3BP-1 is a member of the Ras GTPase-activating protein SH3 domain-binding protein (G3BP) family, named after its identification as an SH3 domain-binding protein of RasGAP (Parker et al., 1996). G3BPs are a highly conserved family of RNA-binding proteins (RBPs) found across eukaryotes (Tourriere et al., 2001). The *Arabidopsis thaliana* genome (TAIR10) encodes eight G3BP homologs that all share conserved structural features (Abulfaraj et al., 2018).

G3BP proteins contain four distinct domains. The NTF2-like domain, located at the N-terminus, is involved in nucleocytoplasmic transport by interacting with RanGTP at the nuclear pore (Suyama et al., 2000), although this function remains to be fully confirmed (Macara, 2001). Additionally, the NTF2-like domain mediates protein-protein interactions and facilitates dimerization of G3BP, which is important for its structural stability and function (Alam & Kennedy, 2019; Kennedy et al., 2001). The acidic and proline-rich (PxxP) region located in the central part of G3BPs facilitates protein-protein interactions, particularly by serving as a binding site for SH3 domain-containing proteins, which are key components of signal transduction pathways (Alam & Kennedy, 2019). The C-terminal region of G3BPs contains two domains essential for RNA binding and regulation, a RGG box and RNA Recognition Motif (RRM). The RGG box, which is commonly found in RNA-binding proteins, enhances RNA binding, nuclear translocation, and post-transcriptional modifications, and thus further links G3BPs to RNA metabolism (Nichols et al., 2000; Darnell et al., 2001; Alam & Kennedy, 2019). The RRM, defined by the RNP1 and RNP2 motifs, interacts with RNA via a β-sheet platform supported by α-helices, and contributes to RNA stability and post-transcriptional modifications (Nagai et al., 1995; Irvine et al., 2004; Alam & Kennedy, 2019).

Recent studies have started to explore the roles of plant G3BPs in stress responses and immunity. Several AtG3BPs form granule-like structures in response to heat stress, which suggests a conserved function for G3BPs in the dynamics of stress granules (SGs) across eukaryotes (Reuper et al., 2021). Moreover, AtG3BP7 (At5G43960) localizes to SGs and contributes to resistance to viruses by interacting with viral proteins (Krapp et al., 2017). Furthermore, transcriptomic analyses revealed that AtG3BPs are differentially regulated under diverse environmental stresses, including heat, cold, high light, and pathogen infections, which suggests G3BPs exert a broader role in stress adaptation (Abulfaraj et al., 2021).

Recent studies identified AtG3BP1 as a negative regulator of plant immunity, as AtG3BP1 loss-of-function mutants exhibit stomatal closure, increased expression of key defense marker genes, and enhanced resistance to *Pseudomonas syringae pv. tomato* (Pst) (Abulfaraj et al., 2018). However, the functional significance of AtG3BP1 phosphorylation in plant immunity remains unclear. As previously stated, our phosphoproteomic analyses identified AtG3BP1 as a target of MPK3, MPK4, and MPK6, which suggests that MPK-mediated phosphorylation modulates the functional activity of AtG3BP1 (Rayapuram et al., 2021). However, the precise role of AtG3BP1 phosphorylation in plant immunity remains uncharacterized.

Herein, we confirm that AtG3BP1 is phosphorylated by MPK3, MPK4, and MPK6. We further demonstrate that phosphorylation at Ser-257 promotes susceptibility to Pst infection by maintaining stomatal opening, suppressing ROS accumulation, and downregulating PTI marker genes and SA biosynthesis. Additionally, we provide evidence that phosphorylation enhances the stability of AtG3BP1 by preventing its degradation via the proteasome. This study uncovers a novel MAPK-dependent phosphorylation mechanism that regulates RNA-binding proteins involved in plant immunity and provides new insights into the post-translational regulation of plant defense responses and may thus contribute to development of new strategies for engineering disease-resistant crops.

## Results

### AtG3BP1 Is Phosphorylated by MPK3, MPK4, and MPK6 In Vitro

Our previous studies indicated that AtG3BP1 is a target of MAPKs (Rayapuram et al., 2021, Van Bentem et al., 2006). To confirm whether AtG3BP1 is a substrate of the immune-activated MAPKs MPK3, MPK4, and MPK6, *in vitro* kinase assays were performed. We purified His₆-MBP-AtG3BP1 as a ∼93 kDa protein. The elution profile (Figure 1A) confirmed successful protein isolation, with fractions E2, E4, E6, E8, and E10 showing consistent bands. Following desalting, the purified protein remained intact, as indicated by a single band on SDS-PAGE (Figure 1B). Next, purified AtG3BP1 was incubated with recombinant MPK3, MPK4, or MPK6 in the presence of ATP, followed by SDS-PAGE analysis using SimplyBlue™ SafeStain. The band corresponding to G3BP1 incubated with each kinase was cut out, as indicated by the arrows in Figures 1C, 1E, and 1G. LC-MS/MS analysis was performed to validate the phosphorylation, and revealed site-specific phosphorylation at Ser-257 of AtG3BP1 (Figures 1D, 1F, and 1H). The fragmentation patterns, supported by annotated b- and y-ions, validated the phosphorylation event, with Mascot scores of 79.5, 77.2, and 72.9 for MPK3, MPK4, and MPK6 respectively. The Mascot Delta (MD) scores further confirmed the phosphorylation data, with values of 36.6, 35.8, and 25.2 for MPK3, MPK4, and MPK6, respectively.

**Figure 1:**
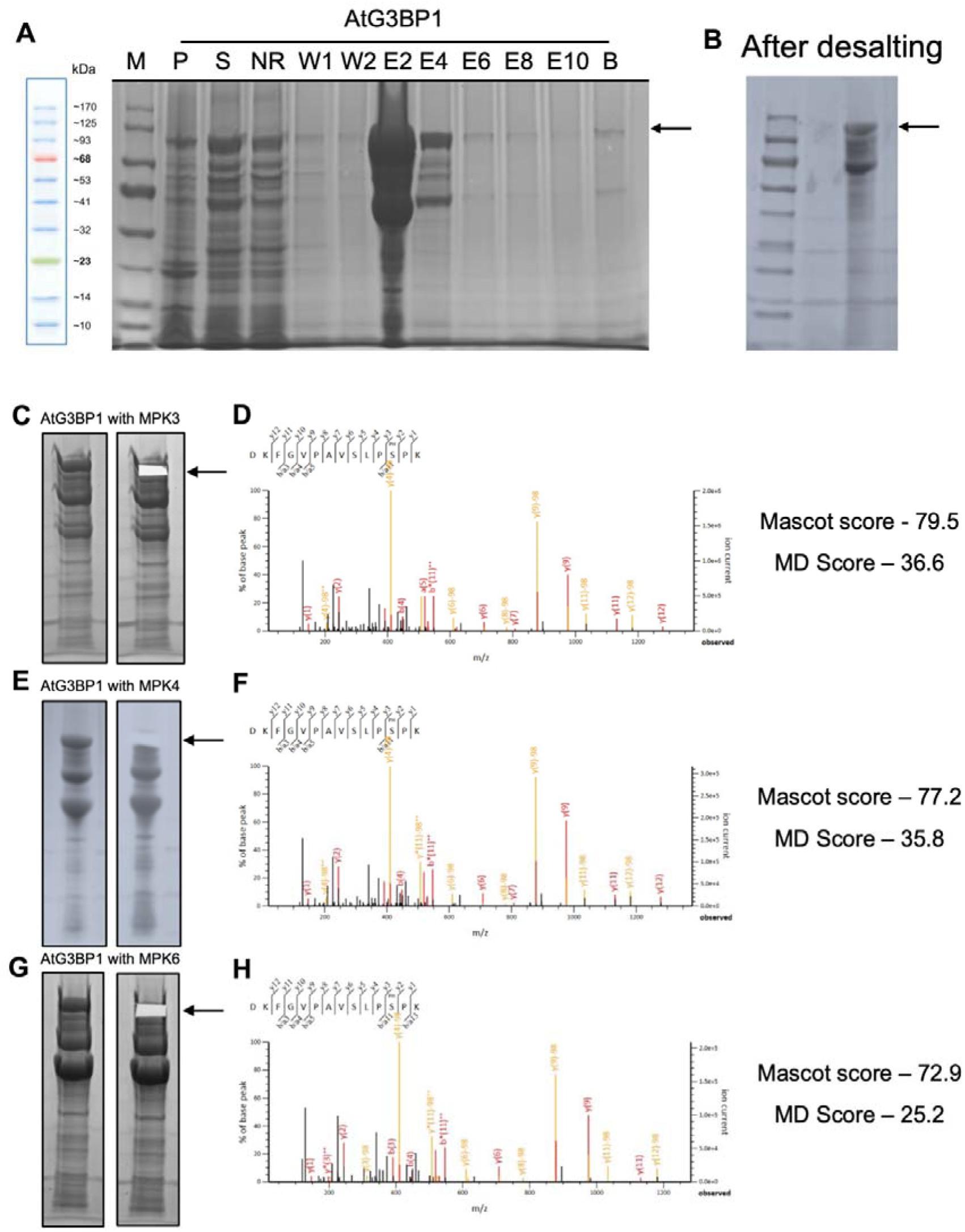
AtG3BP1 is phosphorylated by MPK3, MPK4, and MPK6 in *vitro*. **(A)** Coomassie blue-stained SDS-PAGE gel of the purification steps of His6-MBP-AtG3BP1. Lane M: molecular weight marker; Lane P: pellet fraction; Lane S: soluble fraction; Lane NR: non-retained fraction; Lanes W1 and W2: wash fractions; Lanes E2, E4, E6, E8, E10: eluted fractions; Lane B: proteins bound to Ni-NTA beads. The arrow indicates the purified His6-MBP-AtG3BP1 protein (∼93 kDa). **(B)** Coomassie blue-stained SDS-PAGE gel showing His6-MBP-AtG3BP1 protein after desalting. The arrow indicates the purified AtG3BP1 protein (∼93 kDa). **(C, E, G)** SimplyBlue™ SafeStain SDS-PAGE of AtG3BP1 after phosphorylation by MPK3, MPK4, and MPK6, respectively. Arrows indicate the AtG3BP1 band excised for analysis. **(D, F, H)** LC-MS/MS spectra confirming phosphorylation of AtG3BP1 at Ser-257 by MPK3, MPK4, and MPK6, respectively. Spectra depict peptide fragmentation with annotated b- and y-ions confirming the phosphorylation site. Mascot and Mascot Delta (MD) scores validate the phosphorylation event for MPK3 (79.6, 36.6), MPK4 (77, 35.8), and MPK6 (72, 25.2), respectively.

### Phosphorylation of G3BP1 Does Not Alter Plant Morphology or the Subcellular Localization

To examine the effect of G3BP1 phosphorylation on plant morphology, the growth of wild-type (Col-0), *g3bp1* mutants, and stable *Arabidopsis* lines expressing G3BP1^WT^, G3BP1^A^ (phospho-dead), and G3BP1^D^ (phospho-mimic) under the constitutive ubiquitin promoter was assessed under normal conditions. At 4 weeks, all genotypes displayed similar rosette sizes and leaf morphologies (Figure S1A). At 6 weeks, stem elongation and inflorescence development were comparable across all lines (Figure S1B), indicating that G3BP1 and its phosphorylation does not influence plant growth and morphology overall.

To determine whether phosphorylation of G3BP1 affects its subcellular localization, confocal microscopy was performed on 5-day-old roots stably expressing G3BP1-GFP fusion proteins. GFP fluorescence was predominantly observed in the cytoplasm across all genotypes, with no differences in the patterns of localization observed between G3BP1^WT^, G3BP1^A^, and G3BP1^D^ (Supplementary Figure S2). No nuclear localization of G3BP1-GFP was detected in any genotype, indicating that phosphorylation of G3BP1 does not alter its intracellular distribution.

**Supplementary Figure S1:**
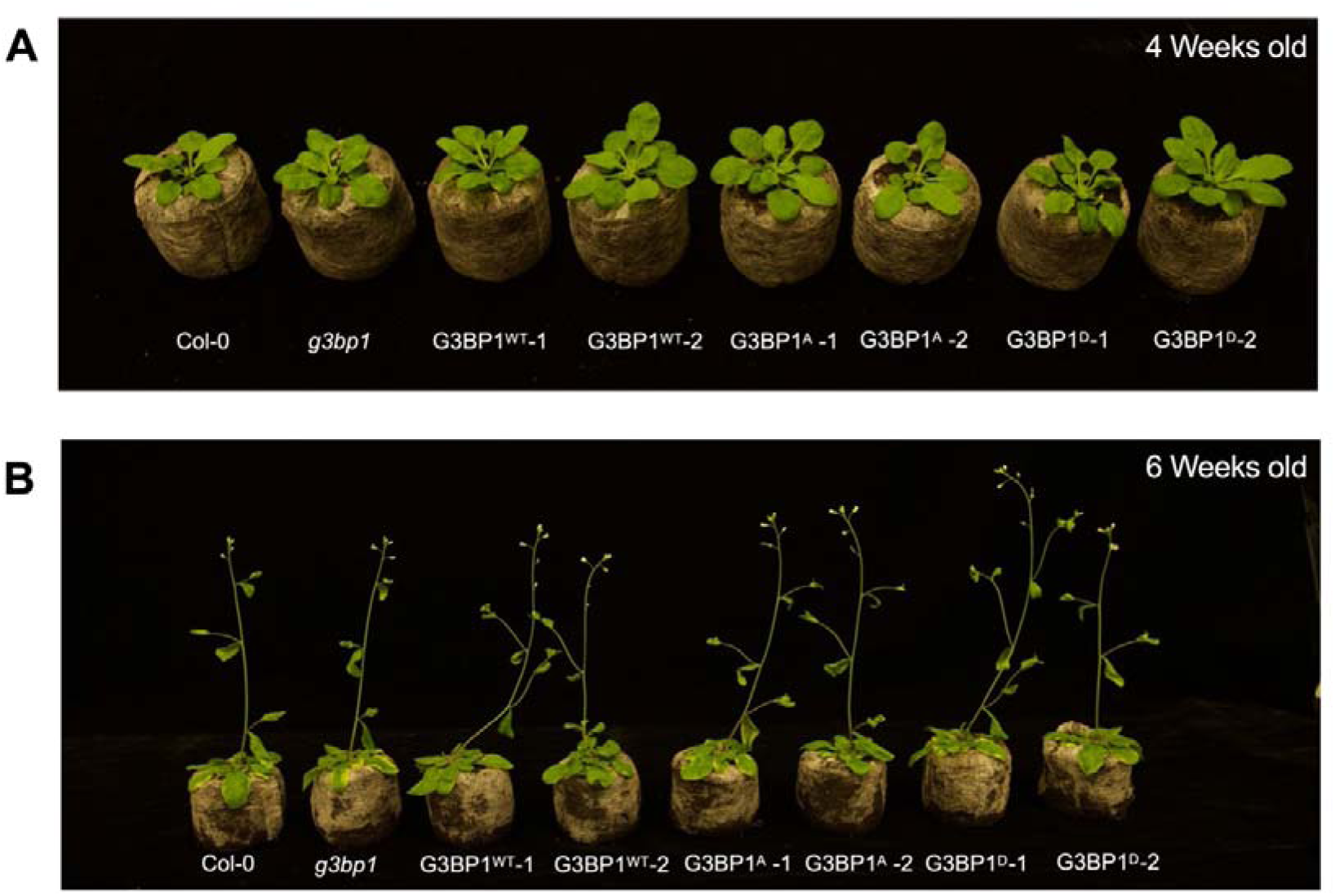
G3BP1 Phosphorylation Does Not Alter Plant Morphology. **(A)** Representative images of 4-week-old wild-type (Col-0), *g3bp1* mutants, and stable *Arabidopsis* lines expressing G3BP1^WT^, G3BP1^A^ (phospho-dead), and G3BP1^D^ (phospho-mimic). **(B)** Representative images of the same genotypes at 6 weeks, showing comparable shoot development and bolting across all lines, indicating that phosphorylation of G3BP1 does not affect plant growth and morphology overall.

**Supplementary Figure S2:**
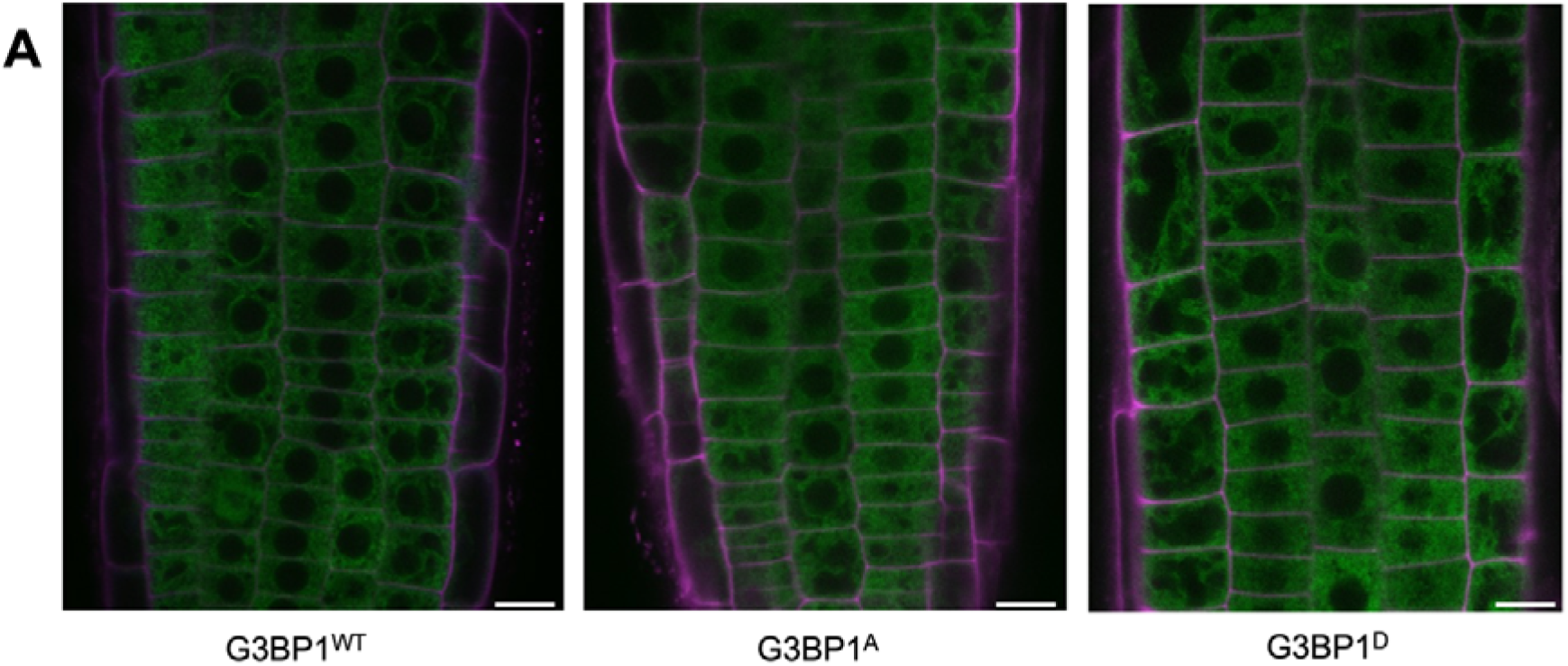
Subcellular localization of G3BP1^WT^-GFP, G3BP1^A^-GFP (phospho-dead), and G3BP1^D^-GFP (phospho-mimic) in *Arabidopsis* root epidermal cells. Confocal laser scanning microscopy images of 5-day-old *Arabidopsis thaliana* roots stably expressing G3BP1^WT^-GFP, G3BP1^A^-GFP (phospho-dead), and G3BP1^D^-GFP (phospho-mimic). GFP fluorescence (green) indicates the localization of G3BP1 fusion proteins, while propidium iodide (PI) staining (magenta) marks the cell walls. Scale bar: 10 µm.

### Phosphorylation of G3BP1 Regulates Plant Resistance to Bacterial Infection and Early Immune Responses

To examine the role of phosphorylation of G3BP1 in plant immunity, pathogen assays were performed using *Pseudomonas syringae pv. tomato* DC3000 (Pst DC3000) in WT (Col-0), *g3bp1* mutants, and stable *Arabidopsis* lines expressing G3BP1^WT^ (wild-type), G3BP1^A^ (phospho-dead), or G3BP1^D^ (phospho-mimic). At both 3 h and 72 h post-infection (hpi), the *g3bp1* mutants and G3BP1^A^ lines exhibited significantly lower bacterial titers compared to WT, whereas G3BP1^WT^ and G3BP1^D^ plants harbored similar bacterial titers as the WT (Figure 2A).

**Figure 2:**
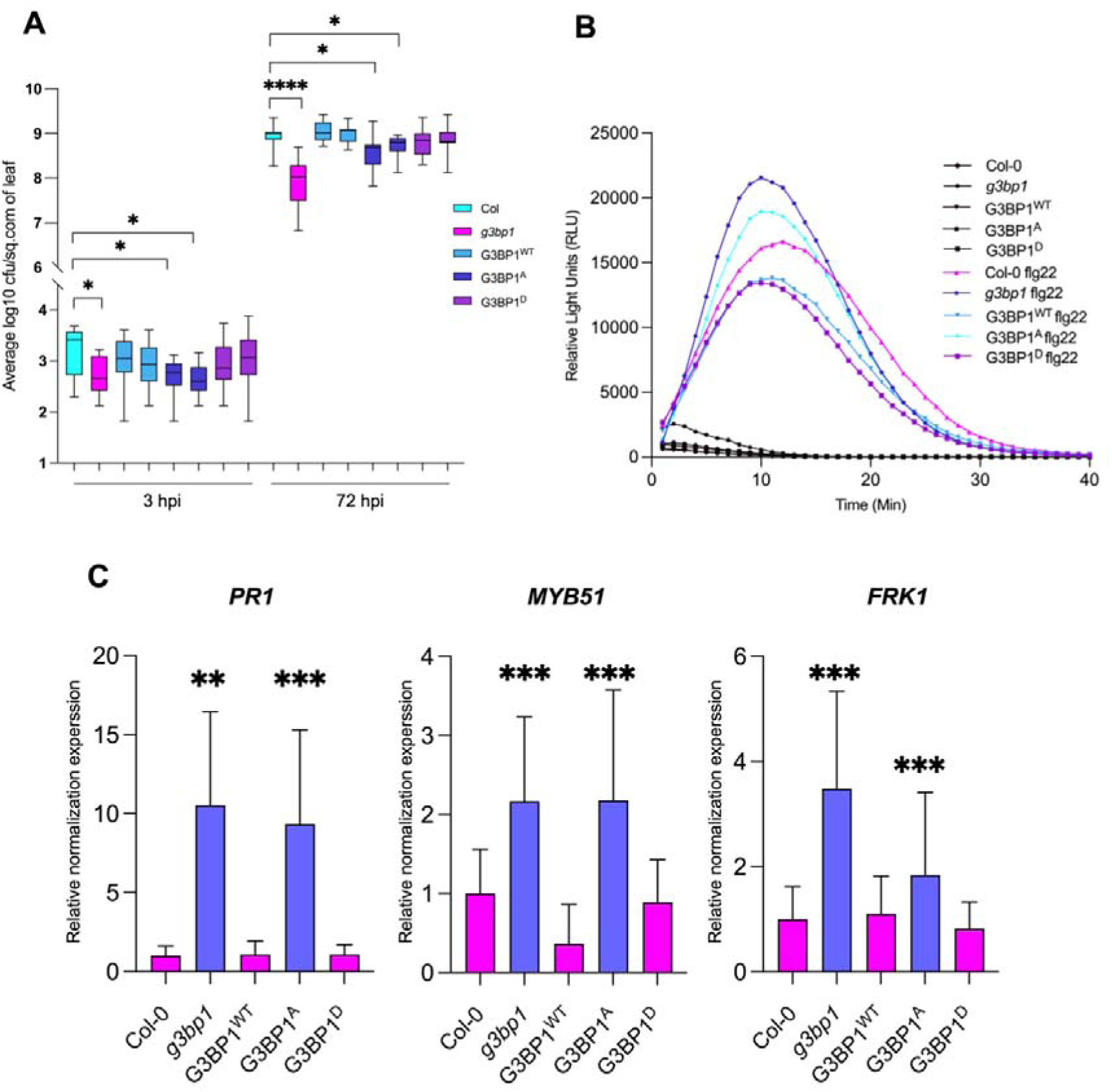
G3BP1 Phosphorylation Regulates Plant Resistance to Bacterial Infection and Modulates Early Immune Responses in *Arabidopsis*. **(A)** Bacterial titers of *Pseudomonas syringae pv. tomato* DC3000 (Pst DC3000) quantified at 3 and 72 hours post-inoculation (hpi) in WT (Col-0), *g3bp1* mutants, and stable *Arabidopsis* lines expressing wild-type G3BP1 (G3BP1^WT^), phospho-dead G3BP1 (G3BP1^A^), and phospho-mimic G3BP1 (G3BP1^D^). The *g3bp1* and G3BP1^A^ plants exhibited reduced bacterial titers compared to WT (Col-0), whereas G3BP1^D^ showed similar susceptibility to WT(Col-0). Data are mean ± SEM of three independent experiments. Statistical significance was determined using the Mann-Whitney U test (*P* ≤ 0.01). **(B)** Reactive oxygen species (ROS) production measured as relative light units (RLU) in response to 1 µM flg22 treatment. Leaf discs from WT (Col-0), *g3bp1* mutants, and stable *Arabidopsis* lines expressing G3BP1^WT^, G3BP1^A^, and G3BP1^D^ were assessed using a luminol-based assay over 40 minutes. The *g3bp1* and G3BP1^A^ plants exhibited significantly enhanced ROS accumulation compared to WT (Col-0) and G3BP1D. Data are mean ± SEM of 12 replicates. **(C)** Expression levels of pattern-triggered immunity (PTI) marker genes (*PR1*, *MYB51, and FRK1*) in WT (Col-0), *g3bp1* mutants, and stable *Arabidopsis* lines expressing G3BP1^WT^, G3BP1^A^, and G3BP1^D^ under mock conditions. Transcript levels were quantified by qRT-PCR, normalized to *UBIQUITIN* and *ACTIN*, and presented relative to WT (Col-0). The *g3bp1* and G3BP1A plants displayed significantly higher expression of PTI marker genes compared to WT (Col-0) and G3BP1^D^. Error bars represent mean ± SD from three independent experiments. Statistical significance was determined using the Student’s t-test. \**P* ≤ 0.05, \*\**P* ≤ 0.01, \*\*\**P* ≤ 0.001, \*\*\*\**P* ≤ 0.0001.

To assess the effects of phosphorylation of G3BP1 on early pathogen-associated molecular pattern (PAMP)-triggered immunity (PTI) responses, reactive oxygen species (ROS) production was measured in leaf discs under mock conditions and after treatment with 1 μM flg22. Upon flg22 treatment, *g3bp1* mutants and G3BP1^A^ plants exhibited significantly higher ROS bursts, generating higher ROS levels than WT (Col-0), G3BP1^WT^, and G3BP1^D^ plants. In contrast, G3BP1^WT^ and G3BP1^D^ plants displayed comparable ROS production levels to WT (Col-0) (Figure 2B).

To further investigate the role of phosphorylation of G3BP1 in immune signaling, the transcript levels of pattern-triggered immunity (PTI) marker genes were analyzed in 14-day-old seedlings under normal growth conditions. The expression levels of Pathogenesis-Related 1 (*PR1*), MYB Domain Protein 51 (*MYB51*), and Flg22-Induced Receptor Kinase 1 (*FRK1*) were significantly upregulated in *g3bp1* mutants and G3BP1^A^ plants compared to WT, G3BP1^WT^, and G3BP1^D^ plants (Figure 2C).

Overall, these results indicate that phosphorylation of G3BP1 modulates plant immunity, with loss of phosphorylation enhancing defense responses, whereas the phospho-mimic G3BP1^D^ behaved similarly to WT plants.

### Phosphorylation of G3BP1 Modulates Stomatal Aperture in *Arabidopsis*

To investigate the role of phosphorylation of G3BP1 in stomatal regulation, stomatal aperture measurements were performed in WT (Col-0), *g3bp1* mutants, and stable *Arabidopsis* lines expressing G3BP1^WT^ (wild-type), G3BP1^A^ (phospho-dead), and G3BP1^D^ (phospho-mimic). Stomatal aperture measurements revealed significant differences between the genotypes (Figure 3A). Under mock conditions, the *g3bp1* mutants and G3BP1^A^ lines exhibited significantly smaller stomatal apertures (median apertures of 0.75 µm and 1.0 µm, respectively). However, compared to WT (Col-0) and G3BP1^WT^ (2.5 µm and 2.0 µm, respectively), G3BP1^D^ exhibited a significantly larger stomatal aperture (∼3.0 µm).

**Figure 3:**
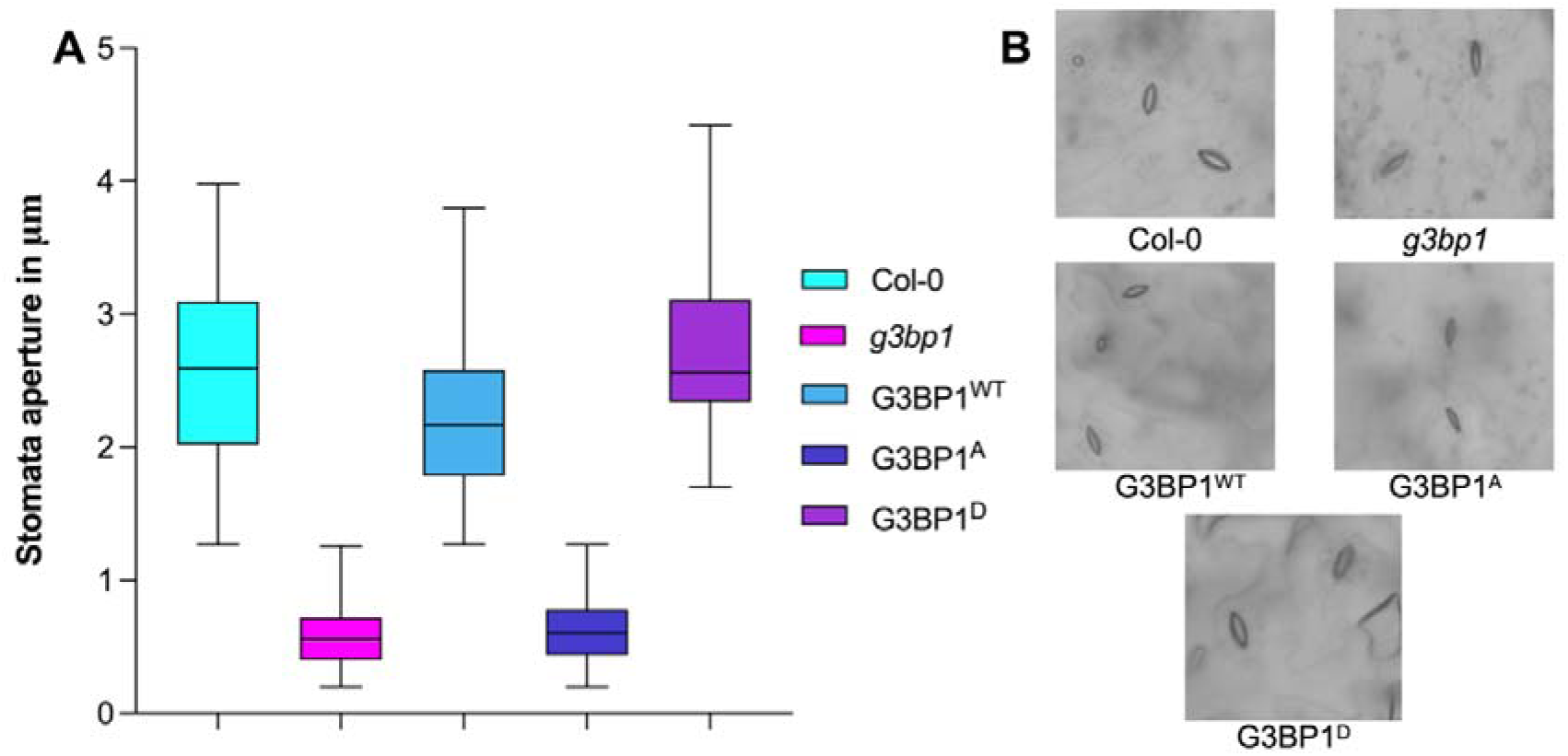
G3BP1 Phosphorylation Modulates Stomatal Aperture in *Arabidopsis*. Stomatal aperture measurements in WT (Col-0), *g3bp1* mutants, and stable Arabidopsis lines expressing G3BP1^WT^ (wild-type), G3BP1^A^ (phospho-dead), and G3BP1^D^ (phospho-mimic). Box plots display median values, interquartile ranges, and whiskers indicate minimum and maximum values. Error bars are the standard error of the mean (SEM). Statistical significance was assessed using one-way ANOVA with Tukey’s multiple comparisons test (*P* < 0.05; \*\**P* < 0.001). **(B)** Representative images of stomatal morphology under mock conditions in WT (Col-0), *g3bp1*, G3BP1^WT^, G3BP1^A^, and G3BP1^D^ plants.

Representative images of stomatal morphology further confirmed the differences in stomatal behavior among genotypes (Figure 3B). WT (Col-0) and G3BP1^WT^ plants exhibited moderately open stomata under mock conditions, while *g3bp1* and G3BP1^A^ lines displayed nearly closed stomata. In contrast, G3BP1^D^ plants showed visibly enlarged stomatal pores, consistent with the quantitative measurements. Overall, these results indicate that phosphorylation of G3BP1 promotes stomatal opening, while loss of phosphorylation leads to constitutive stomatal closure, highlighting its role in modulating pre-invasive immunity.

### Phosphorylation of AtG3BP1 Regulates Stomatal Immunity via SA Signaling

To investigate the effects of phosphorylation of AtG3BP1 on salicylic acid (SA)-mediated defense pathways, we quantified the expression levels of SA biosynthesis, accumulation, and signaling genes in WT (Col-0), *g3bp1* mutants, and stable *Arabidopsis* lines expressing G3BP1^A^ (phospho-dead) and G3BP1^D^ (phospho-mimic) mutant versions of G3BP1.

Quantitative RT-PCR analysis revealed that the SA biosynthesis genes SAR Deficient 1 (*SARD1*), Isochorismate Synthase 1 (*ICS1*), and Calmodulin-Binding Protein 60g (*CBP60g*) were significantly upregulated in *g3bp1* and G3BP1^A^ lines compared to WT (Col-0), while G3BP1^D^ plants exhibited reduced expression of these genes (Figure 4A). Specifically, *SARD1* expression was nearly two-fold higher in *g3bp1* and G3BP1^A^ plants compared to WT (Col-0), but was significantly reduced in G3BP1^D^ plants relative to WT (Col-0). Similarly, *ICS1*, the key enzyme responsible for SA biosynthesis, exhibited the same trend, showing increased expression in *g3bp1* and G3BP1^A^ plants and reduced expression in G3BP1^D^ plants relative to WT (Col-0).

**Figure 4:**
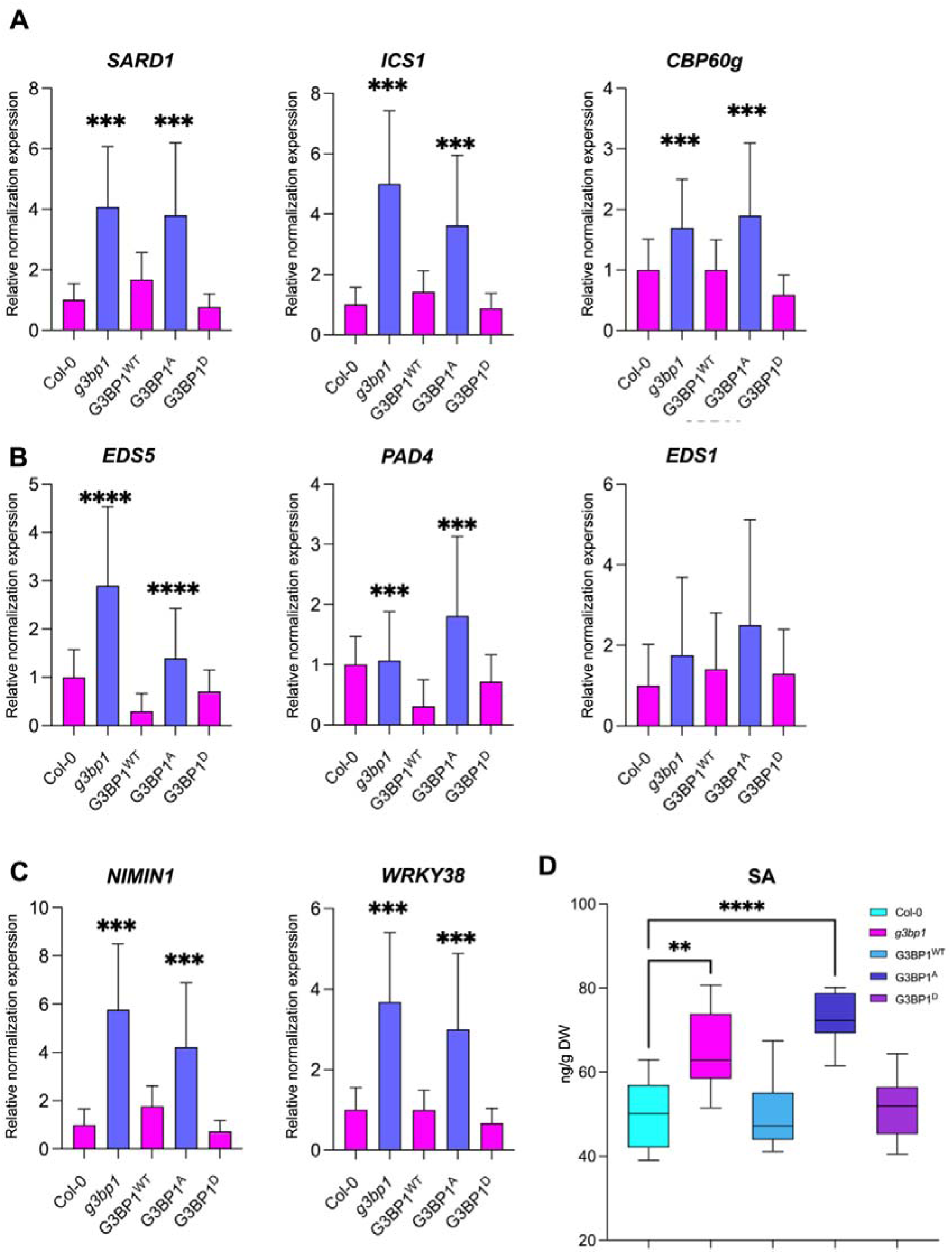
Endogenous Accumulation of Salicylic Acid (SA) in g3bp1 and Phospho-Dead G3BP1A Mutants Correlates with Stomatal Closure and Immune Defense. **(A)** Expression of SA biosynthesis-related genes (*SARD1*, *ICS1*, *CBP60g*) in WT (Col-0), *g3bp1*, G3BP1^WT^, G3BP1^A^ (phospho-dead), and G3BP1^D^ (phospho-mimic) plants. **(B)** Expression of SA accumulation-related genes (*EDS5*, *PAD4*, *EDS1*) in WT (Col-0), g3bp1, G3BP1WT, G3BP1A, and G3BP1D plants. **(C)** Expression of SA signaling-related genes (*NIMIN1*, *WRKY38*) in WT (Col-0), *g3bp1*, G3BP1^WT^, G3BP1^A^ (phospho-dead), and G3BP1^D^ (phospho-mimic) plants. **(D)** Endogenous SA levels in 4-week-old WT (Col-0), *g3bp1*, G3BP1^WT^, G3BP1^A^ (phospho-dead), and G3BP1^D^ (phospho-mimic) plants. Data are shown as box plots with median values, and statistical significance was determined using the Student’s t-test. \**P* ≤ 0.05, \*\**P* ≤ 0.01, \*\*\**P* ≤ 0.001, \*\*\*\**P* ≤ 0.0001.

Genes involved in SA accumulation and transport, specifically Enhanced Disease Susceptibility 5 (*EDS5*), Phytoalexin Deficient 4 (*PAD4*), and Enhanced Disease Susceptibility 1 (*EDS1*), were also significantly upregulated in *g3bp1* and G3BP1^A^ plants compared to WT (Col-0), whereas G3BP1^D^ plants exhibited significantly lower expression of these genes compared to WT (Col-0) (Figure 4B). Notably, *EDS5* expression was nearly three-fold higher in *g3bp1* and G3BP1^A^ plants relative to WT (Col-0), while G3BP1^D^ plants exhibited significantly lower *EDS5* expression compared to WT (Col-0). A similar pattern was observed for *PAD4*, a critical regulator of SA accumulation, which further indicates phosphorylation of G3BP1 negatively regulates SA-mediated defense responses.

The expression levels of NIM1 Interacting 1 (*NIMIN1*) and WRKY DNA-Binding Protein 38 (*WRKY38*), which regulate SA-dependent immune signaling, were significantly upregulated in *g3bp1* and G3BP1^A^ plants compared to WT (Col-0), while G3BP1^D^ plants displayed significantly lower transcript levels compared to WT (Col-0) (Figure 4C). *WRKY38* expression was nearly four-fold higher in *g3bp1* and G3BP1^A^ plants relative to WT (Col-0), whereas G3BP1^D^ plants exhibited significantly lower *WRKY38* expression compared to WT (Col-0). These results suggest that phosphorylation of G3BP1 negatively regulates SA-mediated immune signaling.

To confirm the impact of phosphorylation of G3BP1 on SA levels, the endogenous content of SA was measured in 4-week-old plants (Figure 4D). Consistent with the gene expression patterns, the levels of SA were significantly higher in *g3bp1* and G3BP1^A^ plants compared to WT (Col-0), while G3BP1^D^ plants exhibited significantly lower SA levels relative to WT (Col-0). Specifically, *g3bp1* and G3BP1^A^ plants accumulated nearly twice as much SA as WT (Col-0), whereas G3BP1^D^ plants showed significantly lower SA accumulation relative to WT (Col-0). These findings confirm that G3BP1 phosphorylation suppresses SA biosynthesis and accumulation, and further support the role of (G3BP1^A^) as a negative regulator of SA-mediated immunity.

### Phosphorylation Regulates the Stability of G3BP1 Protein

To determine the impact of G3BP1 phosphorylation on protein stability, we analyzed the levels of G3BP1-GFP fusion protein in stable *Arabidopsis* lines expressing G3BP1^WT^, G3BP1^A^ (phospho-dead), and G3BP1^D^ (phospho-mimic) using immunoblotting. WT (Col-0) was included as a negative control. Total protein was extracted from plants treated with or without the proteasome inhibitor MG132. In the absence of MG132, G3BP1A-GFP protein levels were significantly reduced compared to G3BP1^WT^ and G3BP1^D^, suggesting that the absence of phosphorylation promotes proteasomal degradation of G3BP1 (Figure 5, top panel).

**Figure 5:**
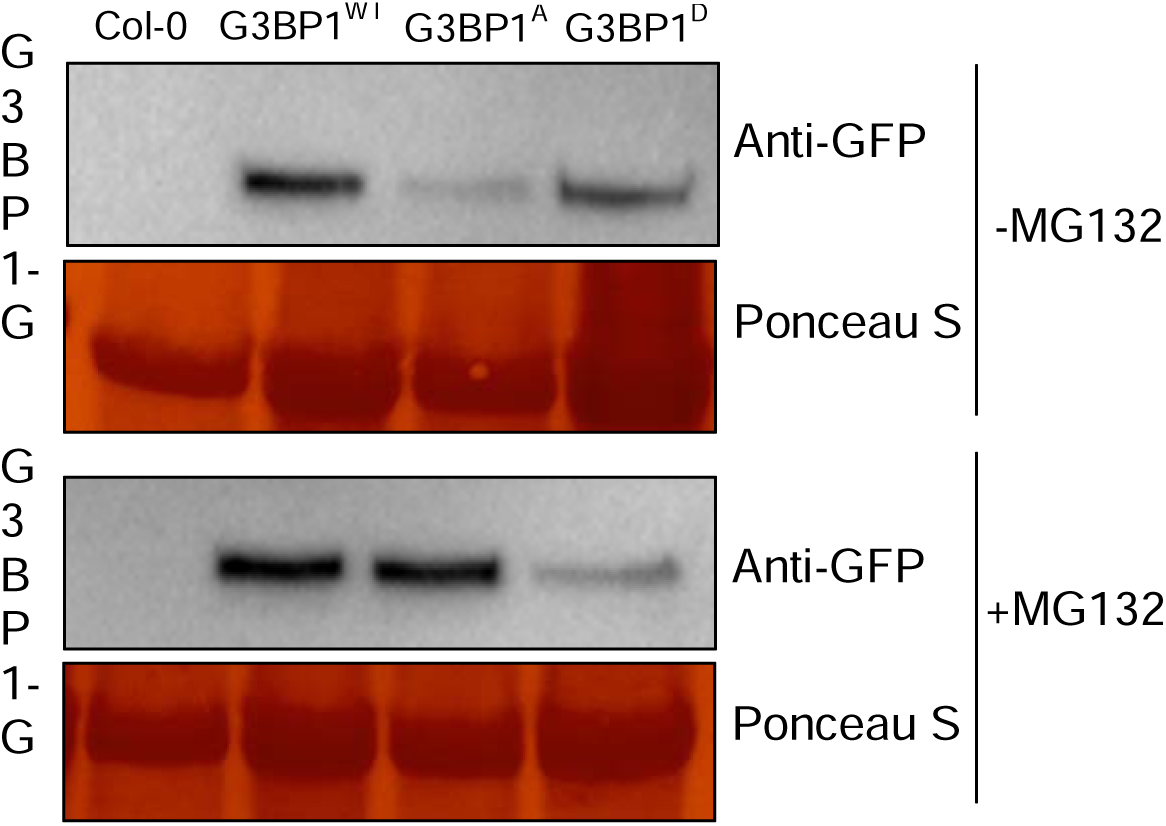
Phosphorylation of G3BP1at Ser275 Phosphosite Affects G3BP1 Protein Stability. Immunoblot analysis of G3BP1-GFP in stable *Arabidopsis* lines expressing G3BP1^WT^, G3BP1^A^ (phospho-dead), or G3BP1^D^ (phospho-mimic). WT (Col-0) was included as a negative control. Total protein was extracted from plants treated with or without 50 μM MG132 for 2 hours. G3BP1-GFP protein levels were detected by immunoblotting using an anti-GFP antibody, and Ponceau S staining was used as a loading control.

To confirm that degradation of G3BP1A-GFP occurs via the proteasome, plants were treated with 50 μM MG132 for 2 hours, followed by protein extraction and immunoblot analysis. MG132 treatment restored G3BP1A-GFP protein levels in the G3BP1A-expressing plants to levels comparable to G3BP1WT and G3BP1D (Figure 5, middle panel), confirming that G3BP1A is degraded through a proteasome-dependent mechanism. Ponceau S staining confirmed equal protein loading across all samples.

## Discussion

Mitogen-activated protein kinases (MAPKs) are essential components of plant immune signaling pathways that transmit extracellular signals to downstream effectors and have been shown to regulate stress responses (Rodriguez et al., 2010). Our findings validate that AtG3BP1, an RNA-binding protein, is a direct phosphorylation target of MPK3, MPK4, and MPK6, revealing a role for in post-transcriptional immune regulation. We demonstrate that phosphorylation of AtG3BP1at Ser-257 modulates plant immunity by promoting susceptibility to bacterial infection, suppressing accumulation of ROS, and downregulating SA biosynthesis. These findings provide novel insights into the role of post-translational modifications in plant defense and establish AtG3BP1 as a key immune signaling component.

The identification of AtG3BP1 as a MAPK substrate expands our understanding of immune signaling through post-transcriptional mechanisms. MAPK cascades can phosphorylate a variety of transcription factors, cytoskeletal proteins, and hormone regulators to control immune responses (Bigeard et al., 2015). Our study highlights the existence of an additional layer of immune control, in that MAPKs regulate RNA metabolism via phosphorylation of AtG3BP1. These findings align with other studies in *Arabidopsis* that have revealed widespread MAPK-mediated phosphorylation of RNA-associated proteins, reinforcing the significance of this regulatory mechanism (van Bentem et al., 2006; Nakagami et al., 2006).

Bacterial growth assays demonstrated that phosphorylation of AtG3BP1 promotes susceptibility to *Pseudomonas syringae pv. tomato* (Pst DC3000). The *g3bp1* mutant and phospho-dead (G3BP1^A^) plants exhibited reduced bacterial titers at 72 hours post-inoculation (hpi), whereas phospho-mimic (G3BP1^D^) plants harbored bacterial titers comparable to WT (Col-0). These findings are consistent with previous reports that AtG3BP1 functions as a negative regulator of SA-dependent immunity, but only in its phosphorylated state (Abulfaraj et al., 2018). MAPK cascades are also known to regulate SA biosynthesis and immune-related gene expression (Bigeard et al., 2015), and our findings suggest that phosphorylation-dependent regulation of AtG3BP1 contributes to this process.

ROS burst assays revealed that *g3bp1* mutants and G3BP1^A^ plants exhibited significantly increased ROS accumulation upon flg22 treatment, whereas G3BP1D plants had reduced ROS levels. ROS are critical signaling molecules involved in plant defense that can amplify immune responses and directly inhibit pathogen growth (Frei dit Frey et al., 2014). The increased ROS levels observed in non-phosphorylated AtG3BP1 plants suggest that phosphorylation of AtG3BP1 suppresses accumulation of ROS and potentially prevents oxidative damage during pathogen defense. MAPK cascades have been shown to modulate ROS homeostasis through phosphorylation of defense-related proteins (Pitzschke et al., 2009).

AtG3BP1 phosphorylation also modulates stomatal immunity, a crucial pre-invasive defense mechanism. Stomatal closure restricts the entry of pathogens into plant tissues, and MAPKs are known to regulate guard cell signaling in response to pathogen attack (Melotto et al., 2006). The *g3bp1* and G3BP1^A^ plants exhibited constitutive stomatal closure, whereas G3BP1^D^ plants maintained larger stomatal apertures. This phenotype is consistent with studies showing that AtG3BP1 mutants exhibit SA-dependent stomatal closure and resistance to coronatine (COR)-induced reopening (Abulfaraj et al., 2018). Since accumulation of SA promotes stomatal closure, and MAPK cascades integrate SA signaling into immune responses (Berriri et al., 2012), our findings suggest that phosphorylation of AtG3BP1 may fine-tune stomatal immunity and play a role in balancing pathogen defense and physiological function.

Our data further indicate that AtG3BP1 phosphorylation negatively regulates SA biosynthesis and signaling. The transcript levels of SA biosynthesis genes (*ICS1*, *SARD1*, and *CBP60g*) and SA accumulation genes (*EDS5*, *PAD4*, and *EDS1*) were significantly increased in *g3bp1* and G3BP1^A^ plants, whereas G3BP1D plants displayed reduced expression of these genes. These results align with previous findings that MAPKs regulate SA-mediated immunity through phosphorylation of transcription factors that control SA biosynthesis (Bigeard et al., 2015). Given that AtG3BP1 mutants accumulate SA (Abulfaraj et al., 2018), our study supports a model in which AtG3BP1 phosphorylation integrates MAPK signaling with SA biosynthetic pathways, which would thereby limit SA-associated defense responses.

Another key finding of this study is that phosphorylation influences the protein stability of AtG3BP1. Immunoblot analysis revealed that G3BP1^A^ protein levels were significantly lower compared to wild-type and G3BP1WT, whereas G3BP1D exhibited greater stability. Moreover, MG132 treatment restored G3BP1^A^ levels, confirming that its degradation is proteasome-dependent. Given that AtG3BP1 was expressed under the control of the ubiquitin promoter in our lines, the observed differences in protein stability are due to post-translational regulation rather than differences in transcriptional control. Our results suggest that phosphorylation enhances G3BP1 stability, potentially by modulating its susceptibility to proteasome-mediated degradation. However, whether phosphorylation directly prevents ubiquitination or alters the protein turnover rate of AtG3BP1 requires further investigation.

Overall, our study establishes AtG3BP1 as a novel MAPK substrate that regulates multiple facets of plant immunity. Phosphorylation at Ser-257 promotes bacterial susceptibility, suppresses ROS homeostasis, inhibits SA biosynthesis, and regulates stomatal aperture, which positions AtG3BP1 as a key immune regulator. In addition, the phosphorylation-dependent stabilization of AtG3BP1 indicates post-translational modifications control protein turnover of AtG3BP1 and immune signaling. Future research is needed to explore whether phosphorylation of AtG3BP1 influences RNA-binding activity, its interactions with stress granules, or directly regulates immune-related mRNAs. Additionally, identification of the upstream stimuli that activate MPK3, MPK4, and MPK6 during infection will help to further elucidate the signaling pathways that govern the function of AtG3BP1. Given its critical role in fine-tuning immune responses, AtG3BP1 presents a promising target for engineering crops with enhanced disease resistance.

## Materials and Methods

### Plant Materials and Growth Conditions

*Arabidopsis thaliana* ecotype Columbia (Col-0) (NASC ID: N1092) was used as the wild-type (WT) plant material, along with the T-DNA insertion mutant *g3bp1*(SAIL_1153_H01) obtained from the Nottingham *Arabidopsis* Seed Centre (NASC).

Seeds were surface-sterilized by vortexing in 1 mL of 70% ethanol containing 0.01% Triton X-100 for 10 minutes at room temperature, followed by three washes with 100% ethanol, dried on sterile Whatman paper in a laminar flow hood, and stratified at 4°C on 1/2 Murashige and Skoog (MS) medium for at least 2 days. The 1/2 MS medium contained Murashige and Skoog basal salts with minimal organics, 0.05% MES hydrate, and 0.5% agar (Sigma A4675, St. Louis, MO, USA), adjusted to pH 5.7 with KOH.

Seedlings were grown in a growth chamber under controlled environmental conditions of 23°C (day)/22°C (night) with a 16 h light/8 h dark photoperiod. For long-term growth, plants were transferred to Jiffy pots and maintained in Percival growth chambers under short-day conditions (8 h light/16 h dark, 60% humidity) at 23°C (day) / 22°C (night) for four weeks.

### *In Vitro* Kinase Assays

*in vitro* kinase assays were performed in a 20 μL reaction volume containing purified recombinant G3BP1 protein and constitutively active MAPKs at a 1:10 kinase-to-substrate ratio, along with 20 mM Tris-HCl (pH 7.5), 10 mM MgCl₂, 5 mM EGTA, 1 mM DTT, and 50 μM ATP. Reactions were incubated at room temperature (RT) for 30 minutes and terminated by the addition of SDS sample buffer, followed by denaturation at 95°C for 10 minutes.

Proteins were resolved by SDS-PAGE and visualized using SimplyBlue™ SafeStain (Novex, Cat. No. LC6065). Protein bands of interest were excised, destained using acetonitrile (ACN) and 100 mM ammonium bicarbonate (NH₄HCO₃) (four washes, 15 min each), and subjected to reduction with 10 mM Tris(2-carboxyethyl) phosphine (TCEP, Sigma, Cat. No. C-4706) at 37°C for 1 hour. Alkylation was performed using 20 mM S-methyl methanethiosulfonate (MMTS, Sigma, Cat. No. 64306) at RT for 30 minutes.

Gel pieces were digested overnight at 37°C with porcine trypsin (Promega), and peptide extraction was performed using ACN and 1% formic acid. Peptides were desalted using C18 ZipTip® columns (Millipore, Cat. No. ZTC18S096) and analyzed by LC-MS/MS. Data were processed using the Mascot server and searched against the TAIR10 database to identify phosphorylation sites.

### Generation of Phosphorylation Lines: Site-Directed Mutagenesis and Generation of Phospho-Mimic and Phospho-Dead Mutants

Site-directed mutagenesis of G3BP1 cDNA was employed to generate phospho-dead and phospho-mimic versions of G3BP1. PCR reactions were carried out using the Phusion High-Fidelity DNA Polymerase (New England Biolabs) and primers designed to introduce specific mutations at serine phosphorylation sites (e.g., S-to-A for phospho-dead and S-to-D for phospho-mimic variants). A DNA plasmid template (5–20 ng) was used as the starting material for each reaction.

Post-PCR, Dpn1 restriction enzyme (Promega) was utilized to selectively digest the parental methylated DNA. The digestion mixture, consisting of 5 μL of purified PCR product and 1 μL of Dpn1 (10 U/μL), was incubated at 37°C for 1 hour. The digested products were then transformed into *E. coli* by heat-shock transformation. Transformants were screened on selective media, and colonies were picked for sequencing analysis to confirm the introduction of the desired mutations.

Confirmed plasmid constructs containing the phospho-dead (S-to-A) and phospho-mimic (S-to-D) mutations were subsequently transformed into *Agrobacterium tumefaciens*. These constructs were introduced into *Arabidopsis thaliana* using the floral dip method. Plants were selected on selective media, and lines expressing the G3BP1 phospho-dead and phospho-mimic variants were identified through three generations of selection and used for subsequent experiments.

### Pathogen Assays

*Arabidopsis* plants (WT, *g3bp1* mutants, and lines expressing G3BP1^WT^, G3BP1A, and G3BP1^D^) were grown for four weeks under short-day conditions (8 hours light/16 hours dark, 22°C). *Pseudomonas syringae pv. tomato* DC3000 (Pst DC3000) cultures were prepared by streaking glycerol stocks streaked onto NYGA plates supplemented with rifampicin (50 mg/L) and incubation at 28°C for 48 hours. For inoculations, bacterial cultures were adjusted to OD_600_ = 0.2 in 10 mM MgCl_2_ containing 0.04% (v/v) Silwet L-77.

Spray inoculations were performed by spraying the bacterial suspension evenly onto the aerial parts of the plants until thoroughly wetted. The inoculated plants were covered with a plastic dome to maintain high humidity and incubated in a growth chamber under short-day conditions. Disease symptoms were evaluated at 3 and 72 hours post-infection (hpi). Leaf discs were collected from three leaves per plant from 10 individual plants per genotype for each biological replicate. Bacteria were extracted by grinding the leaf discs in 10 mM MgCl_2_ containing 0.04% (*v/v*) Silwet L-77.

The bacterial homogenates were serially diluted 10-fold and 10 μL of each dilution was plated on LB agar containing rifampicin (50 mg/L). Plates were incubated at 28°C for 48 hours, and bacterial colonies were counted to determine the colony-forming units (CFU) per cm² of leaf tissue. Data were collected from three biological replicates for each plant genotype.

### ROS Burst Assays

A luminol-based luminescence assay (Huang et al., 2013) was used to measure the production of reactive oxygen species (ROS) in *Arabidopsis* WT (Col-0), *g3bp1* mutants, and lines expressing G3BP1^WT^, G3BP1^A^, and G3BP1^D^. Leaf discs (4 mm in diameter) were excised from four-week-old plants and placed adaxial side-up in 96-well plates (Thermo Fisher, Rochester, NY, USA). Each well contained 150 μL of sterile water, and the plates were incubated overnight at room temperature in the dark. The following day, the water was replaced with 100 μL of reaction solution containing 50 μM luminol (Sigma, St. Louis, MO, USA), 10 μg/mL horseradish peroxidase (HRP; Sigma, St. Louis, MO, USA), and 1 μM flg22 as the microbe-associated molecular pattern (MAMP) elicitor. Water was used as a mock control. Luminescence was recorded using a TECAN Infinite 200 PRO microplate reader at 1-minute intervals for 40 minutes following the addition of the reaction solution. The signal integration time for each measurement was 0.5 seconds. ROS production was quantified as relative light units (RLU), and data were expressed as the mean RLU across three biological replicates, each containing 12 leaf discs per genotype.

### Measurement of Stomatal Aperture

To analyze stomatal responses, epidermal peels from 4-week-old *Arabidopsis* leaves were floated on stomatal opening buffer (10 mM KCl, 50 mM CaCl₂, and 10 mM MES, pH 6.2) under light for 2 hours (Desclos-Theveniau et al., 2012). The peels were then mounted on slides and microscopic images of stomata were captured using a Leica DM5000 confocal microscope. Internal stomatal aperture widths were measured using ImageJ software (Dow & Bergmann, 2014). A minimum of 50 stomata were analyzed per genotype.

### Quantitative RT-PCR (qRT-PCR) Analysis

Total RNA was extracted from 14-day-old *Arabidopsis* seedlings using the RNeasy Plant Mini Kit (Qiagen) according to the manufacturer’s instructions. RNA integrity was assessed with a NanoDrop spectrophotometer (Thermo Fisher Scientific), and 1 μg RNA was used for cDNA synthesis with the SuperScript™ III First-Strand Synthesis SuperMix kit (Invitrogen) and oligo (dT) primers. The cDNA was diluted 10-fold in nuclease-free water. qRT-PCR was performed using SsoAdvanced Universal SYBR Green Supermix (Bio-Rad) in a CFX96 Real-Time System (Bio-Rad). Each reaction contained 2 μL diluted cDNA, 5 μL SYBR Green Supermix, and 2 μL of a 330 nM primer mix, with technical triplicates per sample. Cycling conditions were 95°C for 2 min, followed by 40 cycles of 95°C for 15 s and 60°C for 30 s. Melting curves confirmed the product specificity. The expression levels of *PR1*, *MYB51*, *FRK1*, *ICS1*, *PAD4*, and *WRKY38* were normalized to AT3G18780 (*Actin*) and At4G05320 (*UBQ10*) and presented relative to wild-type (WT) Col-0 controls (set to 1.0). Data were analyzed using Bio-Rad CFX Manager software, with statistical significance determined in GraphPad Prism 9.

### Quantification of Salicylic Acid

Salicylic acid (SA) levels were quantified in leaves of 4-week-old *Arabidopsis thaliana* plants, including Col-0 (wild type), *g3bp1* mutant, G3BP1 ^WT^ (G3BP1 overexpressing wild type), G3BP1^A^ (phospho-dead mutant), and G3BP1^D^ (phospho-mimic mutant). Phytohormone extraction and analysis were performed following a protocol by Trapp et al. (2014). For chromatographic analysis, a Thermo Fisher TQS-Altis Triple Quadrupole Mass Spectrometer was coupled with a Thermo Scientific Vanquish MD HPLC system. Separation was achieved using an UPLC Kinetex C18 column (2.6 μm, 2.1 × 150 mm). The mobile phase consisted of water (A) and acetonitrile (B), with a flow rate of 0.5 mL/min. The gradient elution started at 25% B, increased linearly to 50% B over 2 minutes, then increased to 100% B over the next 6 minutes (by 8 min total). The solvent composition was maintained at 100% B until 11.4 minutes, followed by a rapid decrease to 20% B at 11.5 minutes. The column was re-equilibrated at 20% B until 14 minutes. The column temperature was maintained at 35°C throughout the run.

### Protein Stability Assay

To assess protein stability, G3BP1-GFP levels were analyzed via immunoblotting. Total protein extracts were prepared from 4-week-old plants treated with or without 50 μM MG132 (Sigma-Aldrich) for 2 h. Samples were separated by SDS-PAGE (10%) and transferred onto PVDF membranes (Millipore). Immunoblotting was performed using an anti-GFP (1:5000, Abcam) and a HRP-conjugated secondary antibody (1:15000, Bio-Rad). Signal detection was performed using the ECL detection kit (Thermo Fisher Scientific).

### Statistical Analysis

All experiments were conducted in three independent biological replicates, with each experiment containing at least three technical replicates per condition. Statistical analysis was performed using GraphPad Prism 9. Data were analyzed using one-way ANOVA, followed by Tukey’s post hoc test for multiple comparisons. Statistical significance was set at *p* < 0.05.

## References

1. Abulfaraj, A.A., Mariappan, K.G., Bigeard, J., Manickam, P., Blilou, I., Guo, X., et al. (2018). The Arabidopsis homolog of human G3BP1 is a key regulator of stomatal and apoplastic immunity. The Plant Journal, 94(5), 822–839.

2. Alhoraibi, H., Bigeard, J., Rayapuram, N., Colcombet, J., & Hirt, H. (2019). Plant immunity: The MTI-ETI model and beyond. Current Issues in Molecular Biology, 30, 39–58.

3. Berriri, S., Garcia, A.V., Freschi, L., Gobbato, E., Bourdais, G., Resentini, F., et al. (2012). Constitutively active mitogen-activated protein kinase versions reveal functions of Arabidopsis MPK4 in pathogen defense signaling. The Plant Cell, 24(10), 4281–4293.

4. Bigeard, J., & Hirt, H. (2018). Nuclear signaling of plant MAPKs. Frontiers in Plant Science, 9, 469.

5. Bigeard, J., Colcombet, J., & Hirt, H. (2015). Signaling mechanisms in pattern-triggered immunity (PTI). Molecular Plant, 8(4), 521–539.

6. Dangl, J.L., & Jones, J.D.G. (2001). Plant pathogens and integrated defence responses to infection. Nature, 411, 826–833.

7. Dangl, J.L., Horvath, D.M., & Staskawicz, B.J. (2013). Pivoting the plant immune system from dissection to deployment. Science, 341(6147), 746–751.

8. Desclos-Theveniau, M., Arnaud, D., Huang, T.-Y., Desclos, R., Chien, C.-T., & Zimmerli, L. (2012). The Arabidopsis lectin receptor kinase LecRK-V.5 represses stomatal immunity induced by Pseudomonas syringae pv. tomato DC3000. PLoS Pathogens, 8(2), e1002513.

9. Frei dit Frey, N., Garcia, A.V., Bigeard, J., Zaag, R., Bueso, E., Garmier, M., et al. (2014). Functional analysis of Arabidopsis immune-related MAPKs uncovers a role for MPK3 as a negative regulator of inducible defenses. Genome Biology, 15(6), R87.

10. Gao, M., Liu, J., Bi, D., Zhang, Z., Cheng, F., Chen, S., & Zhang, Y. (2008). MEKK1, MKK1/MKK2 and MPK4 function together in a mitogen-activated protein kinase cascade to regulate innate immunity in plants. Cell Research, 18(12), 1190–1198.

11. Glazebrook, J. (2005). Contrasting mechanisms of defense against biotrophic and necrotrophic pathogens. Annual Review of Phytopathology, 43, 205–227.

12. Huang, T.-Y., Desclos-Theveniau, M., Chien, C.-T., & Zimmerli, L. (2013). Arabidopsis thaliana transgenics overexpressing IBR3 show enhanced susceptibility to the bacterium Pseudomonas syringae. Plant Biology, 15, 832–840.

13. Jones, J.D.G., & Dangl, J.L. (2006). The plant immune system. Nature, 444, 323–329.

14. Kadota, Y., Shirasu, K., & Zipfel, C. (2014). Regulation of the NADPH oxidase RBOHD during plant immunity. Plant & Cell Physiology, 56(8), 1472–1480.

15. Macho, A.P., & Zipfel, C. (2014). Plant PRRs and the activation of innate immune signaling. Molecular Cell, 54(2), 263–272.

16. Meng, X., & Zhang, S. (2013). MAPK cascades in plant disease resistance signaling. Annual Review of Phytopathology, 51, 245–266.

17. Nakagami, H., Soukupová, H., Schikora, A., Zárský, V., & Hirt, H. (2006). A mitogen-activated protein kinase kinase kinase mediates reactive oxygen species homeostasis in Arabidopsis. Journal of Biological Chemistry, 281(50), 38697–38704.

18. Nicaise, V., Roux, M., & Zipfel, C. (2009). Recent advances in PAMP-triggered immunity against bacteria: Pattern recognition receptors watch over and raise the alarm. Plant Physiology, 150(4), 1638–1647.

19. Pitzschke, A., Schikora, A., & Hirt, H. (2009). MAPK cascade signaling networks in plant defense. Current Opinion in Plant Biology, 12(4), 421–426.

20. Rayapuram, N., Jarad, M., Alhoraibi, H.M., Bigeard, J., Abulfaraj, A.A., Völz, R., et al. (2021). Chromatin phosphoproteomics unravels a function for AT-hook motif nuclear localized protein AHL13 in PAMP-triggered immunity. Proceedings of the National Academy of Sciences of the United States of America, 118(3), e2004670118.

21. Rodriguez, M.C., Petersen, M., & Mundy, J. (2010). Mitogen-activated protein kinase signaling in plants. Annual Review of Plant Biology, 61, 621–649.

22. Teige, M., Scheikl, E., Eulgem, T., Dóczi, R., Ichimura, K., Shinozaki, K., et al. (2004). The MKK2 pathway mediates cold and salt stress signaling in Arabidopsis. Molecular Cell, 15(1), 141–152.

23. Tourriere, H., Chebli, K., & Tazi, J. (2003). mRNA degradation machines in eukaryotic cells. Biochimie, 85(8), 609–618.

24. Zhang, Z., Li, Y., Qi, F., Liu, J., & Zhang, Y. (2007). A Pseudomonas syringae effector inactivates MAPKs to suppress plant immunity. Nature, 451(7182), 215–219.

